# SCFAs-induced GLP-1 Secretion Links the Regulation of Gut Microbiome on Hepatic Lipogenesis in Chickens

**DOI:** 10.1101/549857

**Authors:** Jianmei Zhang, Yin shuang Sun, Liqin Zhao, Tiantian Chen, Meina Fan, Hongchao Jiao, Jingpeng Zhao, Xiaojuan Wang, Fuchang Li, Haifang Li, Hai Lin

## Abstract

Chickens represent a specific case in lipid metabolism that liver is the main site of lipid synthesis. As ovipara, their gut microbiota could be strongly influenced by environment and diets after hatching. The aim of this study is to elucidate the linkage of gut microbiota and fat synthesis in broilers. The broilers were subjected to dietary treatments of combined probiotics (*Clostridium butyrate* 4×10^8^ cfu/kg, *Bifidobacterium* 2×10^8^ cfu/kg, *Lactobacillus plantarum* 2×10^8^ cfu/kg and *Lactococcus faecalis* 2×10^8^ cfu/kg, PB) and guar gum (1 g/kg, GG). The result showed that dietary supplementation of PB and GG changed the cecal microbiota diversity, altered short chain fatty acids (SCFAs) contents, and suppressed lipogenesis in liver and abdominal fat tissues. In intestinal epithelial cells (IECs), acetate, propionate, and butyrate upregulated the expression of glucagon-like peptide-1 (GLP-1) via MAPK pathways, especially via the ERK and p38 MAPK pathways. GLP-1 suppressed lipid accumulation in primary hepatocytes with the involvement of AMPK/ACC signaling. In conclusion, the result suggests that SCFAs-induced GLP-1 secretion links the regulation of gut microbiome on hepatic lipogenesis in chickens.

**IMPORTANCE:** Intestinal microbes metabolize SCFAs and stimulate intestinal epithelium L cells to produce GLP-1. Recent evidence showed that GLP-1 reduced fat deposition by reducing appetite and increasing satiety. However, how SCFAs stimulate the secretion of GLP-1 and whether GLP-1 directly affects fat metabolism is not clear. Poultry adipocytes have limited ability to produce fat, and 90% of carcass fat is synthesized in the liver. In addition, large intake of feeds easily leads to fatty liver diseases in chickens. The aim of this study is to investigate how SCFAs mediate secretion of GLP-1 and whether GLP-1 could directly affect hepatic deposition in broiler chickens. The hepatic lipogenesis regulated by the intestinal microbiota of chickens is of great significance to the study of intestinal microbiota and fat deposition in poultry, and this work could provide reference for intestinal microorganism and fat metabolism in mammals and humans.

## Introduction

Gut microbiota plays an important role in the metabolism of the host and altered structure of gut microbiota can affect the energy metabolism (1-4). Gut microbiota can influence both energy balance (weight gain or loss) and energy store (fat mass) by its ability of secreting or altering the metabolites during fermentation (5, 6). The abundance and composition of the gut microbial population are influenced by diet, weight, and overall metabolic state of the host (7). Probiotics and prebiotics are usually used to modulate the gut microbiota towards a beneficial effect on host metabolism (8-11).

The altered gut microbiota changes the end products of fermentation such as short chain fatty acids (SCFAs), which is suggested to be involved in the benefits of microbiota diversity on lipid metabolism (12). SCFAs produced by the gut microbiota in the colon enable the host to gain extra energy, resulting in obesity (13-18). On the other hand, many reports demonstrate that SCFAs could reduce or reverse body weight gains and adiposity (19-22). In obese mice, oral administration of sodium butyrate decreased body weight, via increased energy expenditure and fat oxidation (19). Additionally, in mice fed high-fat diet (HFD), oral administration of acetate, propionate, and butyrate reduced body weight without changing food intake or levels of physical activity (22, 23). SCFAs can serve as signaling molecules via the pathway of specific G protein-coupled receptors GPR43 (FFAR2) and GPR41 (FFAR3) (24, 25). FFAR2 and FFAR3 are demonstrated to be involved in the regulation of lipid and glucose metabolism (26, 27).

Glucagon-like peptide-1 (GLP-1), a neuropeptide derived from the transcription product of the proglucagon gene secreted by the intestinal L cells, is involved in SCFAs-induced metabolism regulation. The activation of FFAR2 and FFAR3 by luminal SCFAs regulates GLP-1 secretion (28, 29). Intra-colonic infusion of propionate elevates GLP-1 levels in portal vein plasma in both rats and mice but not in the FFAR2^−/−^ mice (30). FFAR2-deficient mice fed with a normal diet are obese, whereas mice overexpressing FFAR2 specifically in adipose tissue remain lean even fed a HFD (31).

Different from rodents, the liver is the main site of *de novo* lipid synthesis in chicken (32, 33). The accumulation of triglycerides in liver easily leads to fatty liver syndrome in chickens (34). Moreover, the gut microbiota is mainly affected by rearing environment after hatching, making chicken an interesting model in gut microbiota research. Hence, we hypothesized that SCFAs or GLP-1 might be linkages between gut microbiota and hepatic lipid metabolism in chickens.

The aim of the present study was to demonstrate the link of SCFAs and GLP-1 between gut microbiota and hepatic lipid metabolism. Broilers were fed with a diet supplemented with probiotics or guar gum to alter the animals’ gut microbiota structure and diversity. Then the cecal SCFAs contents and GLP-1 level were measured. The direct effect of SCFAs on GLP-1 secretion and the involved signalings were *in vitro* evaluated with primary intestinal epithelial cells (IECs). The regulation of GLP-1 on hepatic lipid metabolism and the underlying signaling pathway were determined with liraglutide *in vitro* cultured primary hepatocytes.

## Materials and Methods

### Animal experiment

A total of 150 one-day-old broiler (Arbor Acres) chicks were randomly divided into 3 treatment groups, with 5 replicates per group and 10 chicks per replicate, and fed with one of the three diets: high fat basal diet (HFD, feed formula of HFD was listed in Table 1. Control), HFD supplemented with combined probiotics (PB, *Animal bifidobacterium*: 4 × 10^8^ cfu/kg; *Lactobacillus plantarum*: 2 × 10^8^ cfu/kg; *Enterococcus faecalis*: 2 × 10^8^ cfu/kg; *Clostridium butyrate*: 2 × 10^8^ cfu/kg), and HFD supplemented with 1g/kg guar gum (GG). Chicks were reared in an environmentally controlled room. Temperature and lighting were maintained in accordance with commercial conditions. The chickens had free access to feed and water during the whole experimental period. The composition and nutrient levels of the basal diet were listed in supplementary Table 1. All animal experiments were performed in accordance with the “Guidelines for Experimental Animals” of the Ministry of Science and Technology (Beijing, PR China).

**Table 1.** Composition and nutrient levels of the experimental diets (on an air-dry basis)

| Ingredients (%) | 0-3 W | 3-6 W |
| --- | --- | --- |
| Corn | 52.05 | 54.66 |
| Soybean meal | 38.02 | 32.81 |
| Soybean oil | 5.79 | 8.5 |
| Maifan stone powde | 0.99 | 1.13 |
| CaHPO<sub>4</sub> | 2.02 | 1.87 |
| NaCl | 0.32 | 0.29 |
| Lysine | 0.1 | 0.12 |
| Methionine | 0.2 | 0.18 |
| Choline chloride | 0.26 | 0.2 |
| Premix* | 0.25 | 0.25 |
| Calculated nutrient content |  |  |
| Crude protein | 21 | 19 |
| Crude fat | 8 | 10.7 |
| Metabolizable energy, kcal/kg | 3100 | 3300 |
| Calcium | 0.9 | 0.9 |
| Available phosphorus | 0.45 | 0.42 |
| Nacl | 0.35 | 0.32 |
| Digestible lysine | 1.2 | 1.096 |
| Digestible methionine | 0.48 | 0.44 |
| Digestible methionine and cysteine | 0.791 | 0.728 |
| Digestible threonine | 0.828 | 0.744 |
| Tryptophan | 0.264 | 0.235 |
| leucine | 1.547 | 1.432 |
| Isoleucine | 0.841 | 0.75 |
| valine | 0.996 | 0.897 |

Premix provides the following per kg of diet: VA, 8000 IU; VD 3 , 3000 IU; VE, 20 IU; VK, 2 mg; VB 1 , 4 mg; riboflavin, 8 mg; D-pantothenic acid, 11 mg; VB 5 , 40 mg; VB 6 , 4 mg; VB 12 , 0.02 mg; biotin, 0.15 mg; folic acid, 1.0 mg; choline, 700 mg; Fe (as ferrous sulfate), 80 mg; Zn (as zinc sulfate), 75 mg; Mn (as manganese sulfate), 80 mg; Cu (as copper sulfate) 10 mg, I (as potassium iodide), 0.40 mg; and Se (as sodium selenite), 0.30 mg.

At d 21 and d 42 of age, the feed intake and the individual body weight were recorded. The production parameters, including the average daily gain (ADG), average daily feed intake (ADFI) and feed conversion ratio (FCR), were calculated. Three chickens with similar body weights (193.0 ± 3.7 g) were selected from each pen, and 12 chickens were selected per group. The blood samples were drawn from the wing vein using heparinized syringes. Plasma samples were collected and centrifuged at 3,000×g for 10min and were stored at −80°C for further analysis of TG and TCH levels. Chickens were sacrificed immediately after the blood sample collection, the liver and abdominal fat pad were separated, harvested and weighed, liver and abdominal fat percentage was calculated (a proportion of total body weight). Approximately 1 g content of the cecum were collected, 1 g to 2 g tissue samples were collected from the duodenum, jejunum, ileum, cecum, colorectum, liver, and abdominal adipose. All the samples were snap-frozen in liquid nitrogen, and then stored at −80°C for subsequent analysis.

### Primary culture of chicken intestinal epithelial cells

Specific pathogen-free (SPF) chicken eggs were purchased and incubated for 19 days. The chicken embryonic was used for the isolation of primary duodenal intestinal epithelial cells (IECs). The isolation method was previously published and modified to meet our requirements (35-37). Gently extruded the duodenal mucosa and transferred to Hank’s Balanced Salt Solution (HBSS) (Solarbio, Beijing, China), washed with HBSS to remove the mucus and intestinal content until the buffer remained clear. Thereafter, the material was digested with 0.05 mg/ml collagenaseI (MP Biomedicals, Santa Ana, California, USA) at 37 °C under steady agitation for 20min. The material was filtered and larger pieces were discarded, and then centrifuged at 800rpm for 10min. The supernatant was discarded. Cell pellets were washed twice with Hank’s at 800rpm for 10min, cells were resuspended in DMEM-F12 (GIBCO, NewYork, USA) supplement with 10% FBS (Crystalgen, New York, USA), 1×10^7^ cells/well were seeded in 6 well plates and incubated at 37°C and 5% CO_2_. Only wells with over 80% cell confluency after two days of culturing were used for trials.

IECs were rinsed with HBSS for 3 times, starved in non-serum DMEM-F12 medium with 20 mM HEPES (Solarbio, Beijing, China) for 2 h. For dose-response of SCFAs on FFARs, chick intesitinal epithelial cells (IECs) were stimulated with acetate at different concentrations (0, 3, 30, 90 mM) for 24 hours. For time course effect analysis, chick gut epithelial cells were stimulated with acetate (3 mM) for various lengths of time (0, 6, 24, 48 h). After indicated, IECs were starved for 2 h, treated with or without 3mM acetate, 1 mM propionate and 1 mM butyrate respectively for 24h. Then IECs were rinsed with HBSS for 3 times and harvested for subsequent analysis. IECs were treated with SCFAs, with or without the MAPK inhibitors: the ERK-specific inhibitor (UO126,10 μM), JNK inhibitor (SP600125, 20 μM), and p38 inhibitor (SB203580, 10μM), incubated at 37°C in a 5% CO_2_ atmosphere for 2h, collect the cell supernatants and centrifuged for 10min at 3000rpm, total GLP-1 levels in cell supernatants were measured using High Sensitivity GLP-1 Active ELISA-Chemiluminescent (Merck Millipore, EZGLPHS-35K, Germany) according to the reports (38, 39).

### Primary culture of chicken hepatocytes

Chicken embryonic hepatocytes were isolated and cultured following the method previously reported by Hahn et al (40, 41). In brief, hepatocytes were prepared from freshly dissected liver tissue of 17-d-old SPF chick embryo. Liver was collected and washed with salt solution (HBSS; Invitrogen, USA) for three times. During this process, gallbladder, sarcolemma and connective tissue were carefully removed. After that, liver was spliced into pieces (about 1 mm^3^). Collagenase −IV (Sigma, St. Louis, MO, USA) was added to HBSS at *the final concentration of* 0.01 mg/ml, digested at 37°C for 5 min, gently blew it with the disposable pipette for 5 minutes to disperse the cells, filtered and centrifuged for 5 min at 1,000 rpm, the cells were collected and washed with HBSS for 3 times. Density gradient centrifugation was used to separate the hepatocytes from other cells, which was conducted in a layer with 60% Percoll (Sigma, St. Louis, MO, USA). The cell suspension was layered on the Percoll layer and centrifuged for 15 min at 3,000 rpm. Collected the cells and washed it three times with HBSS. The separated cells were counted and seeded at a density of 1×10^7^ cells/mL, cultured in William’s E Medium (GIBCO, NewYork, USA) supplemented with 10% fetal bovine serum (FBS; crystalgen, USA) and 50U/mL penicillin(GIBCO, NewYork, USA), 50µg/mL of streptomycin (GIBCO, NewYork, USA).Cells were incubated in a humidified incubator (Thermo incubator, Forma, USA) at 37°C with 5% CO_2_ for 72 h. The medium was changed at 2 days intervals.

The heptocytes were treated with 100nM chicken GLP-1 (chick GLP-1 sequence: HAEGTYTSDITSYLEGQAAKEFIAWLVNGRG, 42), synthesized by Mimotopes, Hangzhou, China) for 24h in the present of 200μM palmitic acid in the medium. As GLP-1 is easily to be degraded, the heptocytes were treated with 100nM LG (Selleck, Houston, TX, USA) with or without the presence of 100nM exendin(Beyotime, Shanghai, China), the inhibitor of GLP-1R. The cells were harvested for further analysis.

### TG and TCH content measurements

Triglyceride and total cholesterol content was determined with commercial kits (GPO-PAP and CHOD-PAP, Nanjing Jiancheng Biotechnology Institute, Nanjing, China).

### Histological staining

Paraffin-embedded liver and abdominal adipose tissues were sliced into 5 μm sections for hematoxylin and eosin stain (HE, Nanjing Jiancheng Bioengineering Institute, Nanjing, China). The histological features were observed and captured under a light microscope.

### Measurement of SCFA concents

SCFA concentrations were determined using GC–MS assay. Cecal chyme was added to 2 mL of water with phosphoric acid, vortexed and homogenized for 2 min. Then, 2 mL of ether was added to the sample, which was rested for 10 min and centrifuged at 4000 rpm for 20 min at 4 °C. The ether phase was removed after centrifugation, and then the extraction was repeated. The two extracts were combined, volatilized to 2 mL, and injected into the GCMS ISQ LT (Thermo Fisher, USA) and TRACE GCMS ISQ LT (Thermo Finnigan, USA) with the following conditions: column temperature: 100 °C (5 min)–5 °C/min–150 °C (0 min)–30 °C/min–240 °C (30 min); flow rate: 1 mL/min; split ratio: 75:1; carrier gas: helium; column: TG WAX 30 m×0.25 mm×0.25 μm; injector: 240 °C; mass spectrometry: EI source; bombardment voltage: 70 eV; single ion scanning mode: quantitative ions 60,73; ion source temperature: 200 °C; cable temperature: 250 °C; and quantitative analysis method: external standard curve method.

### Cell viability

The cell viability under different treatments was determined by CCK-8 kit (Trans, China) at the wavelength of 450nm.

### Oil Red O Staining

Cells were washed with cold phosphate buffered saline (PBS) and fixed in 10% paraformaldehyde for 30 min. Then the cells were stained for 30 min in a freshly diluted Oil Red O solution (Solarbio, Beijing, China). After rinsed in distilled water, the cells were counterstained for 2 min in Mayer’s Hematoxylin (Sigma-Aldrich, St. Louis, MO, USA).The image of each group was photographed. Subsequently, the stained lipid droplets were extracted with isopropanol for quantification by measuring its absorbance at 490 nm.

### RNA isolation and quantitative real-time PCR analysis

Total RNA from cultured cells or intestinal tract, including the duodenum, jejunum, ileum, cecum and colorectum, liver and abdominal adipose was prepared by the acid phenol method using Trizol reagent (Invitrogen, USA) according to the manufacturer’s instructions. And 1.0 μg total RNA was reverse-transcribed into cDNA using the transcriptor first-strand cDNA synthesis kit (Roche, China). qPCR was conducted using FastStart Universal SYBR Green Master (Rox) reagents (Roche). All the primers were designed by Primer 5, and standard curves and melting curves were performed to ensure the specificity and PCR efficiency. Relative mRNA expression of the target genes was evaluated using the comparative threshold cycle method (2^-ΔΔCT^). Each sample was amplified in triplicate, and GADPH was used as an internal control. Primers used for qRT-PCR were listed in Table 2. Thermal cycling was initiated with an activation step of 30s at 95°C, and this step was followed by 40 cycles of 95°C for 5s and 60°C for 30s.

**Table 2.**
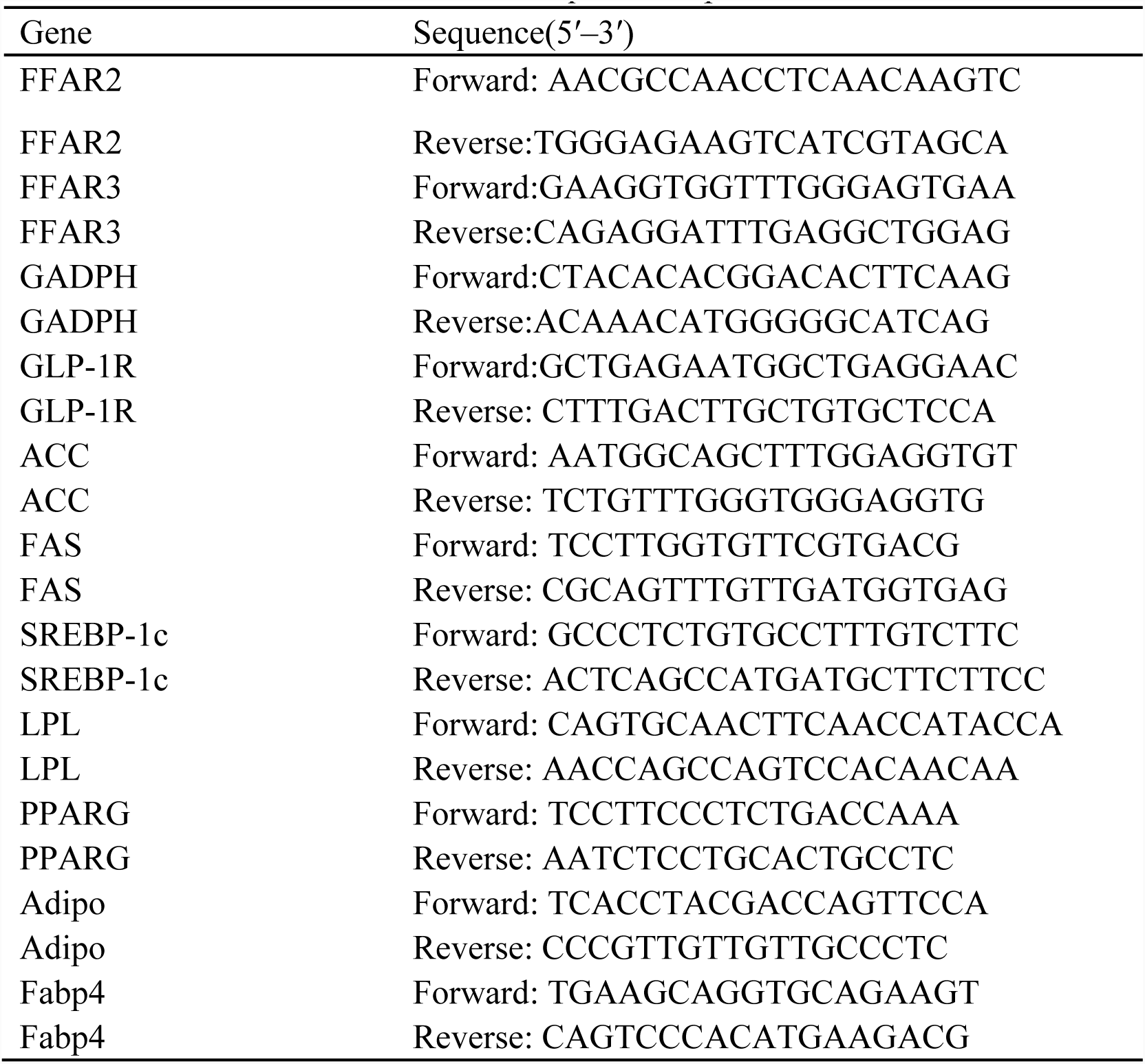
qRT-PCR primers

### Western blot analysis

Total protein extracts from cultured cell lysates or tissue samples were prepared by homogenization in RIPA buffer (1% Nonidet P-40, 0.5% sodium deoxycholate, and 0.1% sodium dodecyl sulfate in PBS) supplemented with protease inhibitor cocktail (Sigma-Aldrich, Oakville, Ontario, Canada) and phosphatase inhibitor cocktail (Fdbio science, China, Alkaline phosphatases, Acid phosphatases, Alkaline phosphatases, PTPs, Ser/Thr phosphatases, PP1, PP2A, PP2B, and PP2C). Cell and tissue homogenates were centrifuged at 12,000×g and 4°C for 10 min. The protein content of the supernatants was determined using the BCA protein assaykit (Beyotime). Total protein (30 μg) was separated by SDS-PAGE and transferred to PVDF membranes (Millipore, Merck, Germany) using a transfer apparatus (BioRad, USA). The membranes were blocked with blocking buffer (Beyotime) at room temperature for 1 h, then incubated with anti-phospho-P38 (#4511T, anti-rabbit, Cell Signaling Technology (CST), USA), anti-P38 (#9212S, anti-rabbit, CST), anti-phospho-JNK (#4668S, anti-rabbit, CST), anti-JNK (#928, anti-rabbit, CST), anti-phospho-ERK (#9101S, anti-rabbit, CST), and anti-ERK (#9102S, anti-rabbit, CST), anti-p-ACC (anti-rabbit, CST), anti-ACC (anti-rabbit, CST), anti-p-AMPK (anti-rabbit, CST) and anti-AMPK (anti-rabbit, CST) primary antibodies overnight at 4°C, followed by incubation with the corresponding horseradish peroxidase-conjugated secondary antibody (Beyotime, Shanghai, China) at 4°C for 4h. Tubulin was used as an internal control for MAPK pathway assay, and β-actin was used as an internal control for other protein expression assays. The protein–antibody complexes were detected with the ECL Plus A and B (Beyotime, Shanghai, China), and the results were quantified using the Fusion FX software (Vilber, France).

### Statistical analysis

The data were expressed as mean±SE and analyzed by one-way ANOVA with SAS software. Differences between means were evaluated using Duncan’s significant difference tests.*p*<0.05 was considered as statistically significant.

## Results

### PB and GG treatments suppressed lipid synthesis and accumulation in the liver and abdominal fat tissue

The GG-chickens had the lowest plasma TG concentration while the control birds had the highest one (P<0.05, Fig. 1a). Compared to the control, the plasma TCH content was decreased by both PB (*p*<0.01) and GG (*p*<0.05) treatments (Fig. 1b). Plasma activity of ALT was decreased in both PB- and GG-chickens compared to control (*p*<0.001) while AST was not changed (*p*>0.05, Fig. 1c,d).

**Figure 1.**
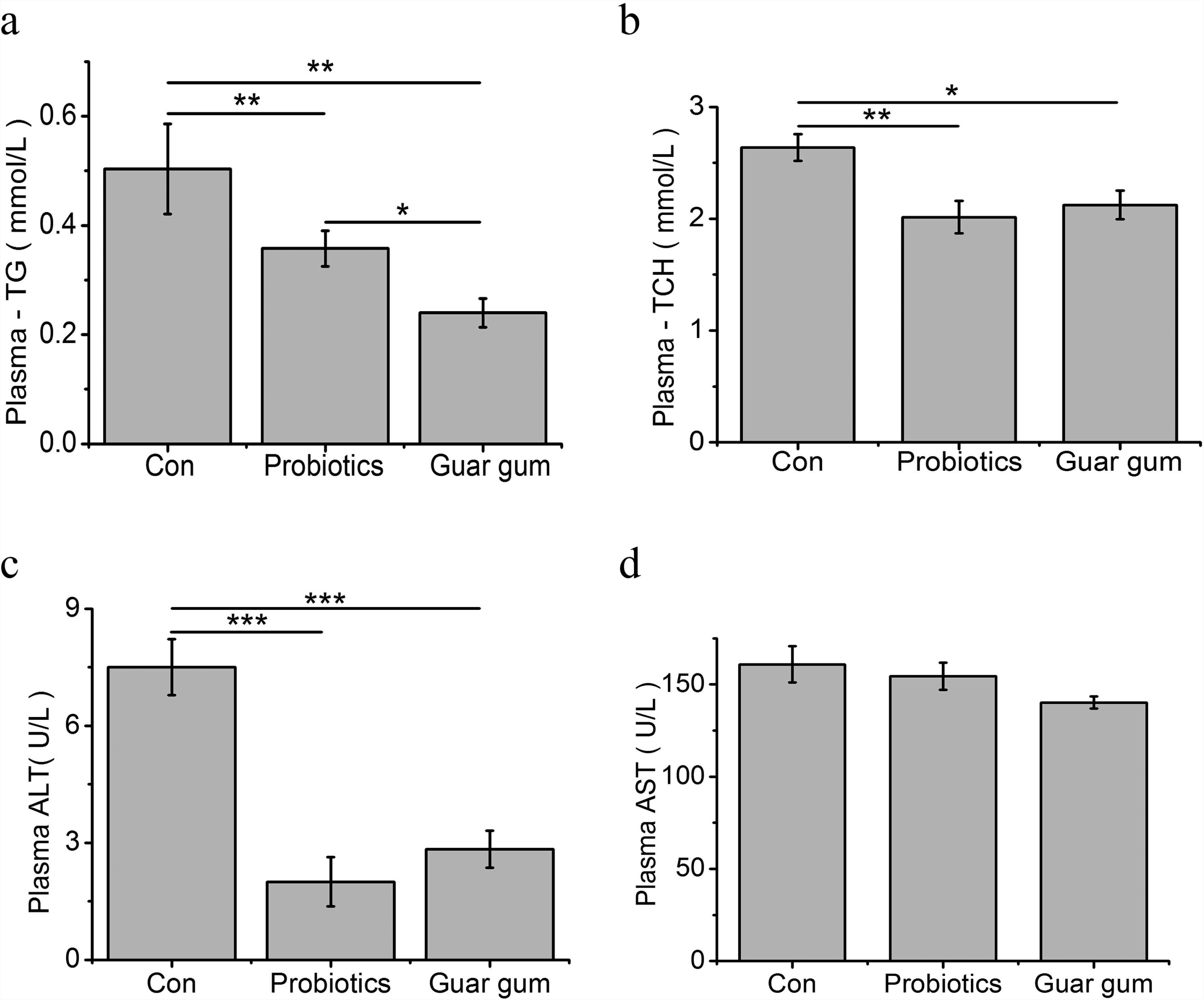
Effect of probiotics and Guar gum supplementations on parameters in plasma of chickens at d 21. (a) TG contents in the plasma; (b) TCH contents in the plasma; (c) ALT activity in the plasma; (d) AST activity in the plasma.SAS analysis followed by T-test (n=8). Data were presented as Mean±SE. \**p* <0.05, \*\**p*<0.01 and \*\*\**p*<0.001.

At d 21, the liver index was significant higher in the PB group compared with control (*p*<0.05, Fig. 2a). The abdominal fat index was significant lower in GG-chickens compared to the control (*p*<0.05, Fig. 2b). At d 42, compared with the control, the liver and abdominal fat index were decreased (*p*<0.05) by PB and GG treatments (Fig. 2a, b). Compared to control, chickens in PB and GG groups showed decreased TG content (*p*<0.05, Fig. 2 c) and alleviated fatty infliltration in liver both 21 and 42 days of age (Fig. 2 d). The hepatic TCH contents, however, was not influenced by either dietary treatment (*p*>0.05, Fig 2 e). In liver, compared to control, the mRNA expression levels of fatty acid synthase (FAS), peroxisome proliferator-activated receptor-γ (PPARG), and sterol regulatory element-binding protein (SREBP)-1c were all down-regulated by GG treatment (*p*<0.05, Fig 2 f). ACC expression was highly suppressed in GG group compared with control (*p*<0.01).

**Figure 2.**
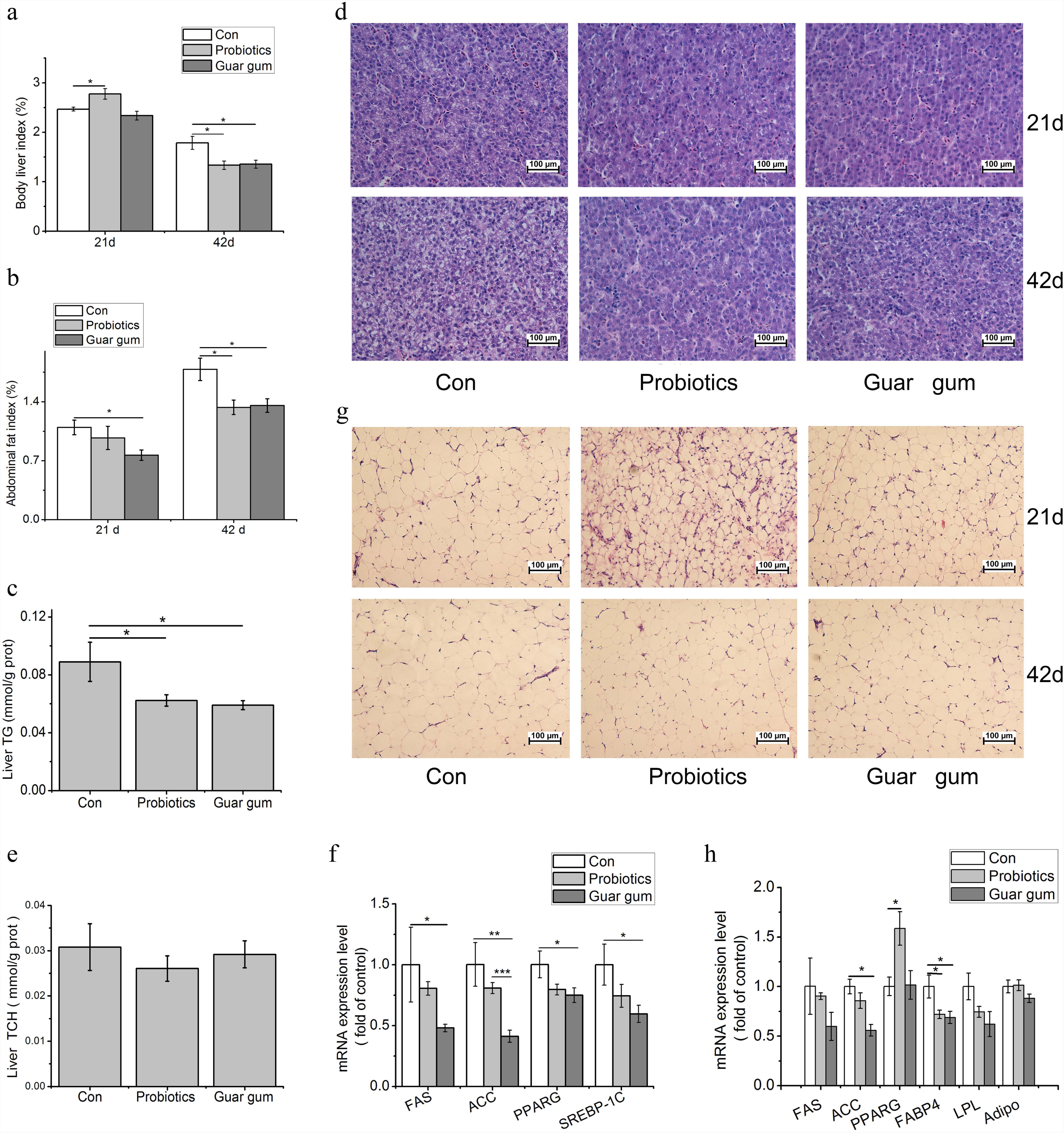
Effects of probiotics and Guar gum supplementation on fat deposition in broilers. (a) liver index at d 21 and d 42 (n=10); (b) Abdominal fat index at d 21 and d 42 (n=10); (c) TG contents in the liver(n=10); (d) H&E staining of the liver slides at d 21 and d 42 (original magnification: ×200, n=6); (e) TCH contents in the liver(n=10); (f) Quantitative RT-PCR analysis of FAS, ACC, PPARG, and SREBP1c at transcript levels in the liver at d 21(n=8); (g) H&E staining of the abdominal fat slides at d 21 and d 42 (original magnification: ×200, n=6); (h) Quantitative RT-PCR analysis of FAS, ACC, PPARG, FABP4, LPL, Adipo (adiponection) at transcript levels in abdominal adipose tissue at d 21(n=8).SAS analysis followed by T-test. Data were presented as Mean±SE. \**p* <0.05, \*\**p*<0.01 and \*\*\**p*<0.001.

**Figure 3.**
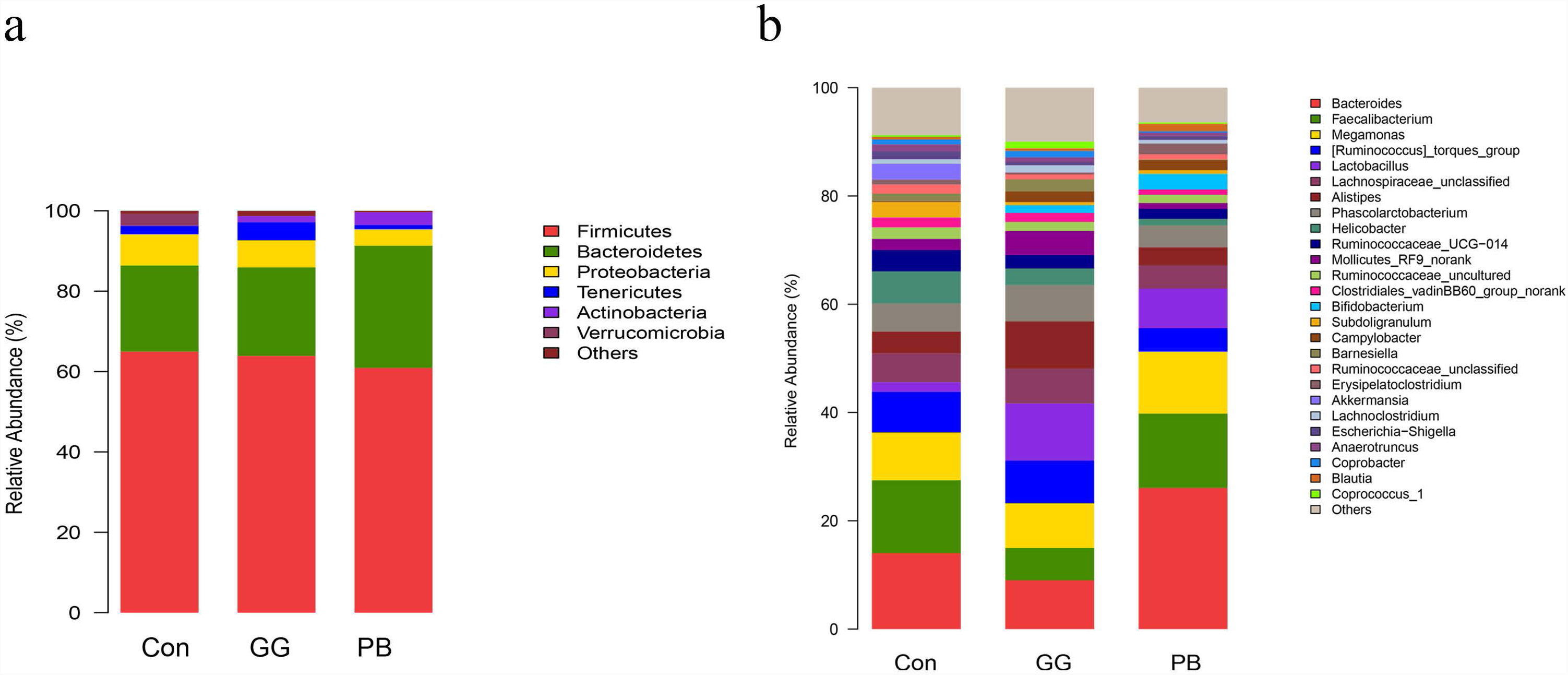
Probiotics and guar gum affect microbiome gut community. Relative abundance of bacteria population in cecal microbiota of groups Control, PB and GG at 21 days of age, sequenced were analyzed by using Illumina MiSeq System. (a) Phylum level; (b) Genus level. (n=8).

In abdominal fat tissue, the average adipocyte size was reduced in abdominal fat padin PB- and GG-chickens compared with control (Fig 2 g). The mRNA levels of ACC and FABP4 in GG-chickens were decreased compared to control (*p*<0.05, Fig 1h). In contrast, PB treatment increased PPARG while decreased FABP4 compared to control (*p*<0.05).

### PB and GG treatments changed the cecal microbiota diversity

Compared to the control group, GG did not change the relative abundances of *Firmicutes* and *Bacteroidetes* at the phylum level, while PB decreased the relative abundance of *Firmicutes* and increased the relative abundance of *Bacteroidetes*. There was a decrease in the relative abundances of *Proteobacteria* in both PB and GG groups. The minor taxonomic groups such as *Actinobacteria* and *Verrucomicrobia* showed the same trends in PB and GG treatments. In line with that at the phylum level, minor taxonomic groups of cecal microbiota in PB and GG groups presented the same trends and showed significant differences with that in control group. Compared with the control, the *Escherichia-Shigella, Anaerotruncus, Akkermansia, Ruminococcaceae_uncultured, Subdoligranulum, Ruminococcaseae* and *Helicobacter* presented much lower abundance in both PB and GG groups, while *Bifidobacterium, Campylobacter* and *Lactobacillus* were significantly higher in PB and GG groups.

### PB and GG treatments are associated with altered cecal SCFAs concentrations

Compared to the control, cecal acetate concentrations were increased in both PB and GG groups (*p*<0.05, Fig. 4 a), while butyrate levels were only increased by GG treatment (*p*<0.05). The propionate content was not influenced by PB or GG treatment (P>0.05). The relative percentage of acetate was not changed by either dietary treatment, whereas the percentage of propionate was decreased in PB treatment, and the percentage of butyrate was increased in GG treatment, compared to control (*p*<0.05, Fig 4 b).

**Figure 4.**
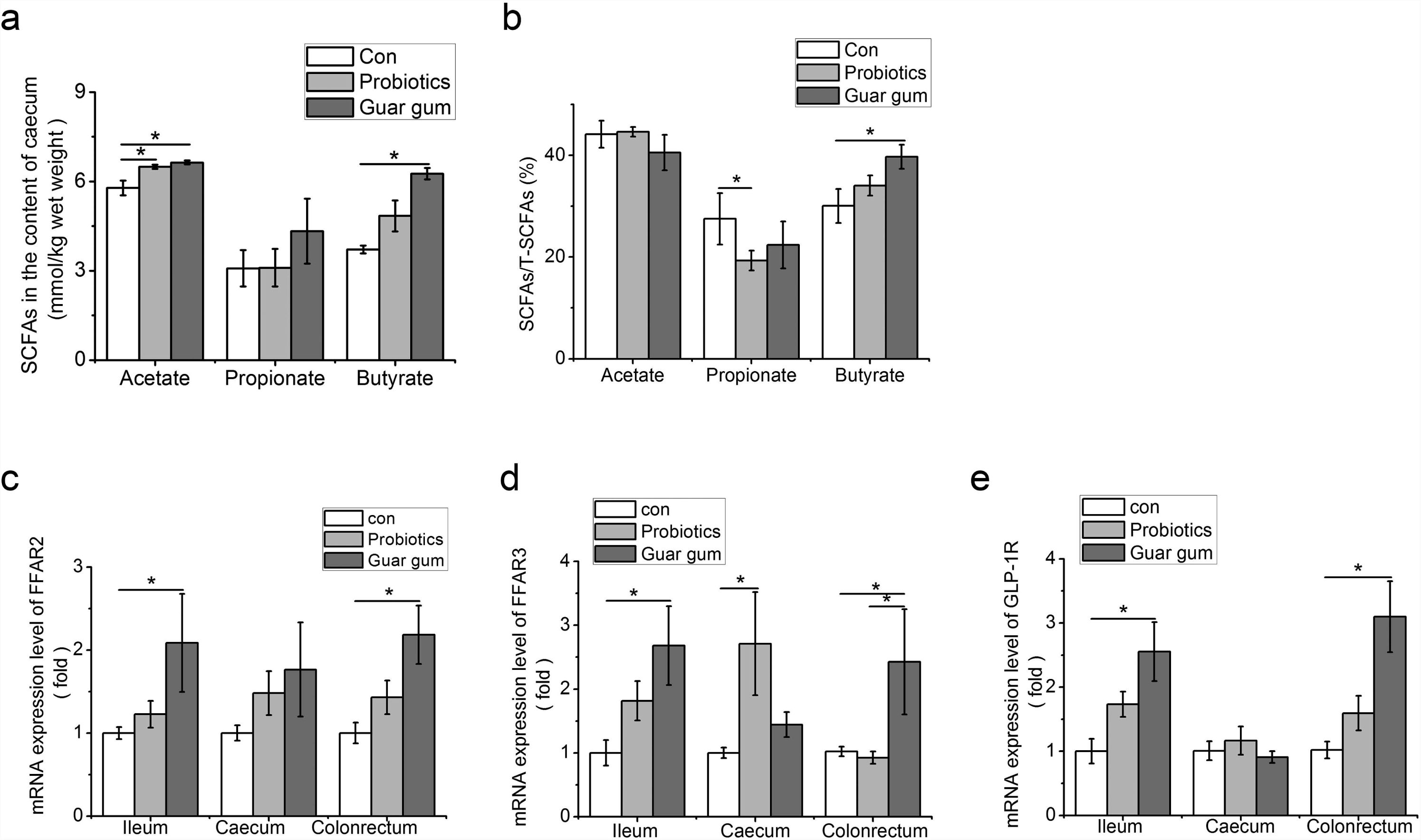
Higher contents of SCFAs increase FFARs and GLP-1R expression in the intestine of both PB and GG groups at 21 days of age. (a) SCFAs contents in the content of caecum (mmol/g wet weight) of HFD chickens in control, PB and GG groups at 21 days of age (n=10 in control and PB groups; n=8 in GG group); (b) SCFAs/ Total SCFA(T-SCFAs) in the content of caecum of HFD chickens in control, PB and GG groups at 21 days of age (n=10 in control and PB groups; n=8 in GG group); (c-e) Effect of probiotics and guar gum on mRNA expression level of FFAR2/3 and GLP-1R in the intestine of HFD chickens at 21 d of age (n=8): (c) FFAR2; (d) FFAR3; (e) GLP-1R.*p<0.05.SAS analysis followed by T-test. Data were presented as Mean±SE, \**p* <0.05.

### PB and GG treatments upregulated the expression of FFARs and GLP-1R

Comparing with control, the mRNA levels of FFAR2 and GLP-1R were upregulated by GG treatment in ileum, ceacum and colonrectum (*p*<0.05), but not in cecum (P>0.05, Fig. 4 c, d, e). Except for FFAR3 mRNA level was increased by PB treatment in ceacum (*p*<0.05) (Fig. 4 d), no significant influence was observed for FFAR2, FFAR3, and GLP-1R mRNA levels in other tissues by PB treatment.

### SCFAs induced GLP-1 secretion via activating MAPK pathways in IECs

Compared to the control, the expression levels of FFAR2, FFAR3, and GLP-1R were significantly upregulated by acetate (*p*<0.05, *p*<0.05, *p*<0.01), propionate (*p*<0.05, *p*<0.01, *p*<0.01), and butyrate (*p*<0.01, *p*<0.05, *p*<0.01; Fig. 5 a, b, c) in IECs. The GLP-1 content in culture medium was increased by acetate (*p*<0.001), propionate(*p*<0.05), and butyrate(*p*<0.05) treatment (Fig 5 d). The phosphorylation levels of ERK and p38 were all significantly increased (*p*<0.001) by acetate, propionate,and butyrate treatment (Fig 5 e, f, g, h). The phosphorylation of JNK was only significantly upregulated by butyrate (P<0.05).

**Figure 5.**
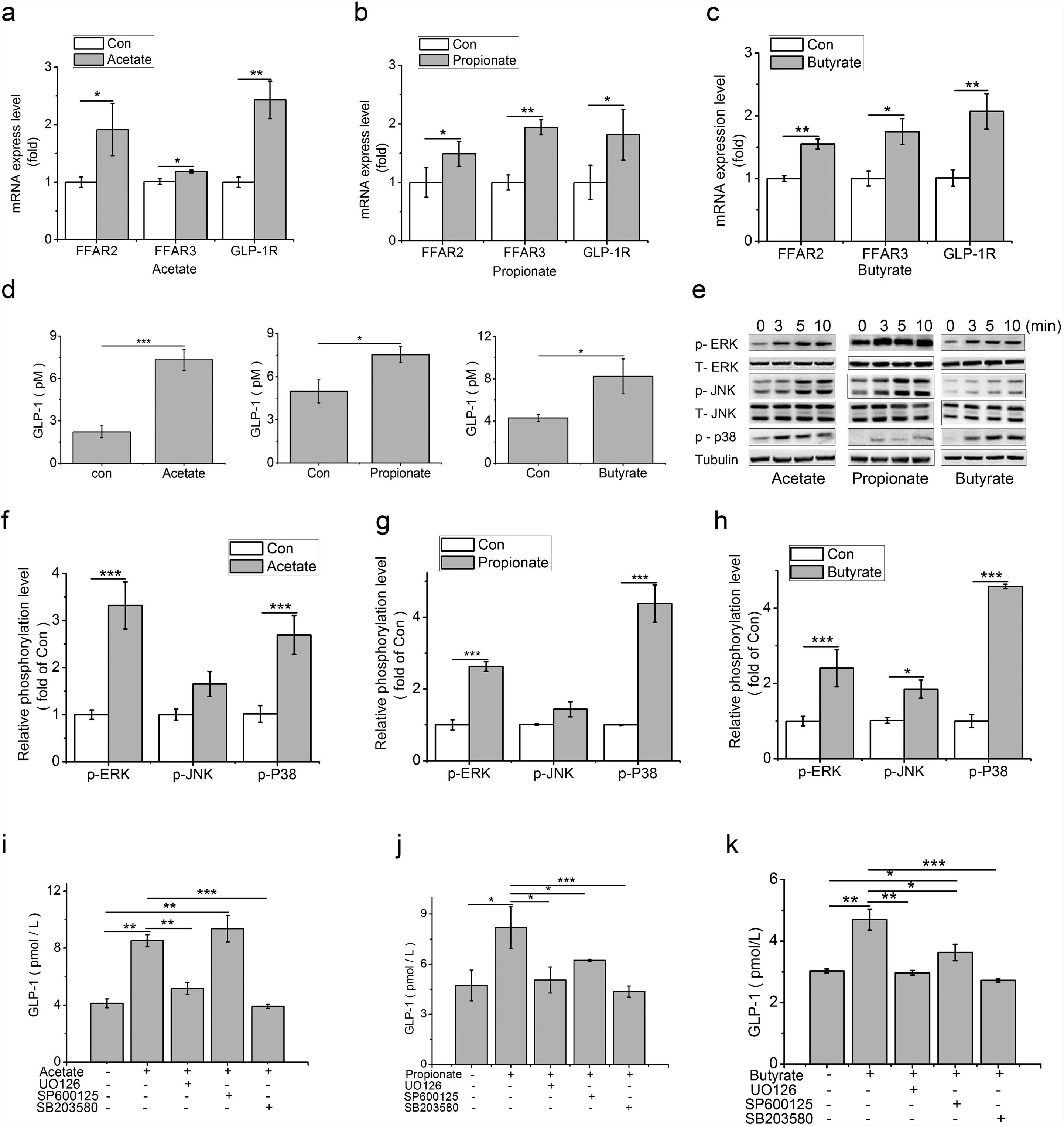
SCFAs stimulate the FFAR2/3 and GLP-1R expressions and mediate GLP-1 secretion in IECs, and SCFAs mediated GLP-1 secretion via MAPKs pathways. (a-c) Transcript levels of FFAR2/3 and GLP-1R were assessed by qPCR in primary IECs after treatments with 3mM acetate, 1mM propionate and 1mM butyrate for 24 hrs (n=8): (a) FFAR2; (b) FFAR3; (c) GLP-1R; (d) GLP-1 concentrations in the culture of the IECs were detected by high sensitive GLP-1 Elisa kit (n=6) after stimulated with 3mM acetate, 1mM propionate and 1mM butyrate for 2 hrs respectively (n=8); (e) SCFAs-mediated activation of ERK, JNK and p38MAPK. IECs were treated with 3mM acetate, 1mM propionate and 1mM butyrate respectively for the indicated times. Intracellular levels of pERK, pJNK, and p-p38 were analyzed by western blotting, relative phosphorylation levels were calculated by p-p38, and pERK/ERK and pJNK/JNK and normalized for control (n=4); (f-h) Quantity of pERK, pJNK and p38MAPK at 5min under the stimulation of SCFAs: (f) Acetate; (g) Propionate; (h) Butyrate; (i-k) MAPK inhibitors block SCFA-induced increase of GLP-1 secretion. IECs were treated with 3mM acetate, 1mM propionate and 1mM butyrate, with or without the ERK-specific inhibitor (UO126,10 μM), JNK inhibitor (SP600125, 20 μM), and p38 inhibitor (SB203580, 10 μM) respectively, GLP-1 concentration was detected by high sensitivity GLP-1 Active ELISA-Chemiluminescent kit (n=6): (i) Acetate; (g) Propionate; (h) Butyrate; Significant comparisons were calculated by SAS with a post T-test. Data were presented as mean ± SE. \**p* <0.05, \*\**p*<0.01 and \*\*\**p*<0.001.

Compared to control, acetate treatment increased GLP-1 concentration (*p*<0.01), which was abolished in the presence of UO126 (*p*<0.01) and SB203580 (*p*<0.001), the inhibitor of ERK and p38, but not influenced by SP600125, the inhibitor of JNK (Fig 5 I). In contrast, the stimulating effect of propionate on GLP-1 secretion (*p*<0.05) was inhibited by UO126 (*p*<0.05), SB203580 (*p*<0.05), and SP600125 (*p*<0.001, Fig 5 J). Similarly, UO126 (*p*<0.01) and SB203580 (*p*<0.001) arrested the stimulating effect of butyrate on GLP-1 secretion (*p*<0.01).Compared to control and butyrate treatment, however, SP600125 treatment increased and decreased, respectively, the GLP-1 secretion (*p*<0.05, Fig 5 K).

### GLP-1 and its analogue liraglutide suppressed lipid accumulation through AMPK/ACC phospharalation in primary hepatocytes

In primary hepatocytes, thelipid accumulation that stained with Oil Red O and quantitated using a spectrophotometer at 500nm and TG contents were decreased by GLP-1 treatment, compared to the control (*p*<0.01, Fig. 6 a,b). Compared to the control, phosphorylation levels of AMPK and ACC were increased by GLP-1 (*p*<0.001, Fig 6 b).

**Figure 6.**
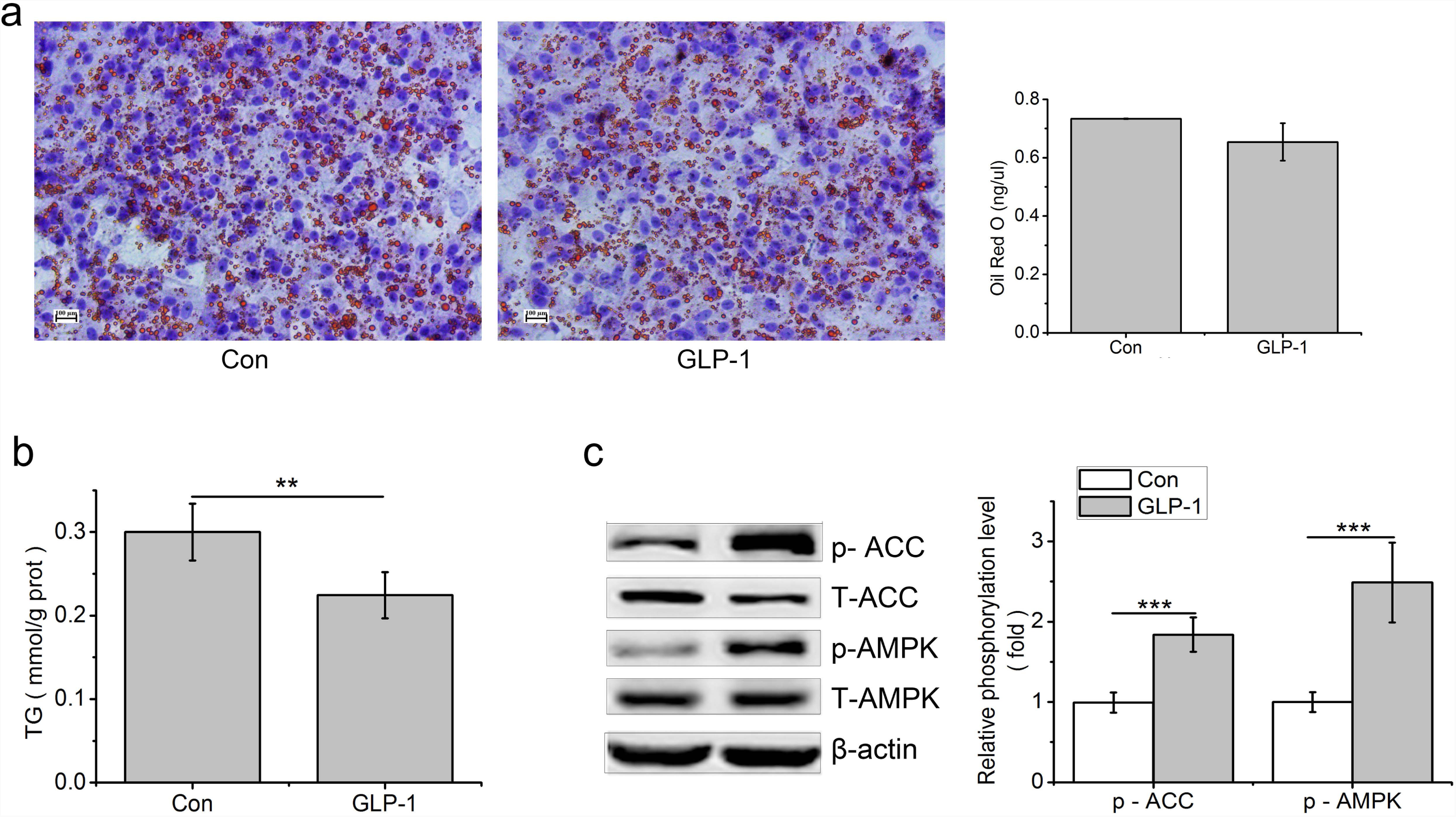
GLP-1 reduces lipid accumulation in the primary hepatocytes. Hepatocytes were stimulated with 100 nM GLP-1 for 24 h, (a) Oil red O staining (original magnification: ×200, n=6); (b) TG content in hepatocytes was measured and normalized to the total protein; (c) GLP-1 increased the phosphorylation of AMPK and ACC. Intracellular levels of pAMPK and pACC were analyzed by western blotting, relative phosphorylation levels were calculated by and the ratio of pAMPK/AMPK and pACC/ACC and normalized for control. Significant comparisons were calculated by SAS with a post T-test. Data were presented as mean ± SE (n= 6). \**p* <0.05, \*\**p*<0.01 and \*\*\**p*<0.001.

Liraglutide treatment decreased the Oil Red O and TG content in the primary hepatocytes, compared to control (*p*<0.01, Fig. 7 a,b). Liraglutide upregulated the transcription level of GLP-1R (*p*<0.01), while downregulated the mRNA levels of PPARG (*p*<0.05), FABP4 (*p*<0.05), LPL (*p*<0.05), and SREBP-1C (*p*<0.001)(Fig. 7 c).

**Figure 7.**
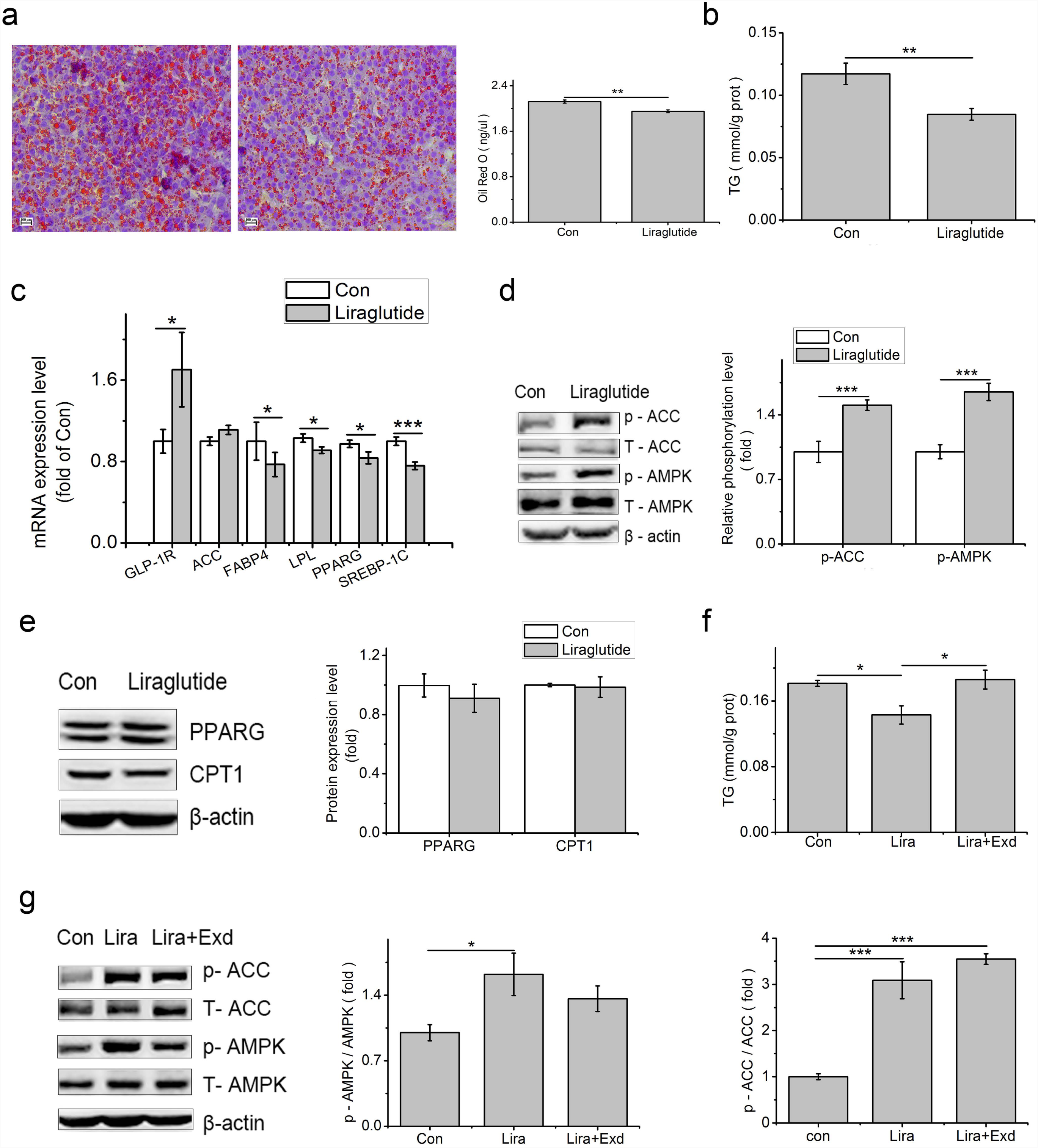
Liraglutide can mimic the effect of GLP-1 on lipid accumulation in the primary hepatocytes. Hepatocytes were stimulated with 100 nM liraglutide for 24 h, (a) Oil red O staining (original magnification: ×200, n=6); (b) TG content in hepatocytes was measured and normalized to the total protein (n=6); (c) Liraglutide reduced transcript expression levels of genes associated with lipid accumulation at primary hepatocytes. mRNA expression of GLP-1R, ACC, FABP4, LPL, PPARG, and SREBP-1C were assessed by qPCR after 24 hrs stimulation with liraglutide (n=6); (d) Liraglutide increased the phosphorylation of AMPK and ACC in primary hepatocytes. Gene expressions were analyzed by western blotting, quantification is shown in the right panel. Relative phosphorylation levels were calculated by the ratio of pAMPK/AMPK and pACC/ACC and normalized for control (n=4); (e) Liraglutide has no effect on the expression of PPARG and CPT1 in primary hepatocytes (n=4). (f) Exendin(9-39)(Exd) recovered TG content Primary hepatocytes; (g) Exendin(9-39) inhibited the phosphorylation of AMK in primary hepatocytes (n=6).Primary hepatocytes were stimulated with liraglutide for 2 h for the detection of protein phosphorylation or for 24 h for TG detection in the presence or absence of 100 nM exendin (9–39). Triglyceride content in hepatocytes was measured and normalized to the total protein in the samples. Gene expressions were analyzed by western blotting, relative phosphorylation levels were calculated by the ratio of pAMPK/AMPK and pACC/ACC and normalized for control. Data were presented as mean± SE; \**p*<0.05, \*\**p*<0.01, \*\*\**p*<0.001.

Compared to control, liraglutide increased the phosphorylated AMPK (*p*<0.001) and ACC (*p*<0.001, Fig. 7 d), but had no influence on PPARG and CPT1 (*p>*0.05, Fig. 7e). In Comparation with control, the decreased TG content caused by liraglutide was restored by exendin (9-39) (Fig. 7 f). In contrast, the phosphorylation level of AMPK in exendin (9-39)+liraglutide treatment had no difference compared with control or liraglutide treatment (*p*<0.05). The phosphorylation level of ACC, however, was not changed by liraglutide (*p*>0.05) (Fig. 7 g).

## Discussion

In the present study, results showed that dietary supplementation of PB and GG changed the cecal microbiota structure and diversity, altered cecal SCFA concentrations, and suppressed fat deposition in liver and abdominal fat tissues. Acetate, propionate, or butyrate could all upregulate the production of GLP-1 via MAPK pathways in IECs. GLP-1 suppressed lipid accumulation in primary hepatocytes with the involvement of AMPK pathway. These findings suggest that SCFAs-induced GLP-1 secretion links the regulation of gut microbiome on hepatic lipogenesis in chickens.

### Altered gut microbiota is associated with the increased SCFAs and decreased lipid accumulation

Gut microbes have been shown to regulate host physiology, metabolism and energy storage (43-45). Altered gut microbial composition have been observed in animals and humans with metabolic syndrome (46, 47). John and Mullin (2016) indicated that obesity accompanies with the increase of *Firmicutes* and decrease of *Bacteroidetes* (48). Though large amounts of correlation research have been reported, the linkage of gut mcrobiome with metabolism still remains largely to be elucidated.

This study indicated that both PB and GG could change the cecal microbiota structure and diversity. In line with previous studies in human and rodents (13, 49), the decreased proportion of phylum *Firmicutes* and increased proportion of phylum *Bacteroidetes* were observed in PB group. At the genus level, the relative abundance of *Lactobacillus, Bifidobacteria* and *Campylobacter* were increased in both PB and GG groups. *Lactobacilli* and *Bifidobacteria*were indicated to produce SCFAs during the fermentation of carbohydrates (50, 51). Meimandipour *et al.* (2010) showed that *Lactobacillus salivarius* and *L. agilis* increased propionate and butyrate contents in cecum of chickens (52). In this study, supplementation with PB and GG also leads to increased ceacal contents of acetate and butyrate, but not with propionate. This is partially in agreement with studies that *Lactobacillus* and *Bifidobacterium* increased the commensal metabolite butyrate (53).

Meanwhile, decreased hepatic and plasma TG and TCH concentrations, and reduced abdominal fat ratio were observed in PB- and GG-treated chickens. Similar effects associated with probiotics and guar gum were observed in mammals. Wang *et al*. (2017) demonstrated that probiotic *Lactobacillus johnsonii* lowered fat deposition by improving lipid metabolism in broilers (54). Guar gum can be digested and produce SCFAs in the hindgut (22, 55), prevent and reverse body weight gain in rodents and humans (56, 57). den Besten *et al*. (2015) showed guar gum protects against HFD-induced obesity via the same signaling cascade as SCFAs (22). Other works demonstrated that increased circulating SCFAs are associated with reduced adipocyte lipolysis and adipogenesis (12, 58). Dietary supplementation of acetate, propionate, and butyrate inhibits lipolysis and *de novo* lipogenesis and protects against HFD-induced obesity (19, 20, 59). One main finding of this study is that liver is also a vital target of gut microbiota regulation on fat metabolism in chickens, in which SCFAs may be linkages.

### SCFAs induce the secretion of GLP-1 via MAPK pathways

Although the intracellular mechanism is not fully understood, luminal SCFAs are expected to stimulate FFAR2 and/or FFAR3 located on the colonic L cells and induce glucagon like peptide-1(GLP-1) release in mammals (28). In the present study, the increased contents of SCFAs were accompanied with higher expression levels of FFARs and GLP-1R in chicken intestines. Hudson *et al*. (2012) showed that FFAR2 and FFAR3 respond to acetate and butyrate at the same level, while FFAR3 was more sensitive to propionate than FFAR2 in mouse (60). A slightly different nature from that in mice, our results showed that both chicken FFAR2 and FFAR3 responded to propionate and butyrate, while FFAR3 was a little more sensitive than FFAR2 in *vitro*. But FFAR2 was much more sensitive to acetate than FFAR3 in primary IECs.

In primary cultured IECs, our results showed that all three SCFAs increased the release of GLP-1. It has been demonstrated that luminal and especially vascular infusion of acetate and butyrate significantly increase colonic GLP-1 secretion, while propionate has no influence on GLP-1 secretion whether administered luminally or vascularly in rat (61). In contrast, although the effect of propionate on GLP-1 secretion is the weakest, it could significantly promote the secretion of GLP-1 in IECs in the present study. Meanwhile, GLP-1 secretion showed a 3-fold increase by acetate and 2-fold increase by butyrate respectively. FFAR2 activation is suggested to predominate over FFAR3 signaling induced by SCFAs with regards to increased gut hormone release (29, 62). Tolhurst *et al*. (2012) reported that SCFAs stimulate GLP-1 secretion via FFAR2 in mixed colonic cultures in mice (29). It suggests that the different GLP-1 secretion may be related to the different activation of FFAR2 under stimulation of acetate, propionate and butyrate. We found that acetate had the strongest stimulating effect on FFAR2, followed by butyrate acid (p<0.01) and propionate (p<0.05) in IECs in this study.

FFAR2 and FFAR3 are G protein coupled receptors (GPCRs). In human and rodents, the coupling of FFAR2 and FFAR3 to ERK1/2 was confirmed under the stimulation of SCFAs in CHO-K1 cells (63-65). Yonezawa et al. (2007) showed that all three SCFAs rapidly and selectively activated p38 MAPK in MCF-7 cells (66). JNK can also be activated by acetate, propionate or butyrate in HEK293 cells, but has rarely been studied due to its lower activation level compared to ERK (65). It is well known that ERK is a major regulator of cell proliferation, whereas JNK and p38MAPK are involved in stress signaling and many inflammatory processes (67). Does MAPK pathway participate in the SCFAs mediated GLP-1 secretion and involve in lipid metabolism? Our present study showed that ERK were stimulated by acetate for 3.5 fold, propionate for 3 fold, and butyrate for 2.5 fold, compared to control. In contrast, significant JNK activation was only detected under butyrate treatment. This result was in agreement with the study that ERK1/2 phosphorylation level mediated by GPR43 and GPR41 were extremely higher with over 3-4 fold to the control, while activation of JNK1/3 was weak (65). The activated p38MAPK was observed in all three SCFAs treatments, which was disagree with the work in parental HEK293 cells, where the activation of p38MAPK was weak under both FFAR2 and FFAR3 stimulators (65). Additionally, in consistent with highly activated ERK and p38MAPK under the stimulation of all three SCFAs, the GLP-1 secretions were reduced to the basal level by inhibitors of ERK1/2 and p38MAPK. In contrast, the acetate mediated GLP-1 secretion is not sensitive under the inhibitor of JNK. These fingdings suggest that ERK and p38MAPK pathways are mainly involved in SCFAs-induced GLP-1 secretion in chickens, which are different from that in mammals.

### GLP-1 and its analog Liraglutide decrease hepatic fat synthesis through phosphorylation of AMPK and ACC

SCFAs have been shown to stimulate the gut hormone GLP-1 release (22), and result in a reduced body weight gain and decreased fat deposition (29, 58, 68). Their main targets are adipose tissue. In this study, we found that both GLP-1 and liraglutide could decrease lipid accumulation in the primary cultured chicken hepatocytes, indicating liver was also a target for GLP-1. Studies demonstrate that treatment with GLP-1 improves liver histology and reduces body weight in animal models of NASH (69, 70). It is suggested that GLP-1 reduces body weight gain and fat deposition by suppressed food intake (68, 71). In ob/ob mice, 60 days of treatment with GLP-1R agonist significantly reduced weight gain and hepatic lipid content, suggesting that GLP-1 has a direct effect on hepatocyte fat metabolism in liver and the GLP-1-treated hepatocytes showed elevated cAMP production as well as reduced mRNA expression of genes associated with fatty acid synthesis (72). Our study showed that GLP-1 and its analog liraglutide significantly increased the phosphorylation of AMPK and ACC (acetyl-CoA carboxylase), whereas had no effect on PPARG and carnitine palmityl transferase I (CPT1) at the protein level in chicken primary hepatocytes. AMPK plays a key role in regulating energy metabolism. Activated AMPK can phosphorylate and inactivate ACC which leads to a decrease in fatty acid synthesis (73, 74), as well as down-regulates transcription factors and enzymes associated with lipid metabolism, such as SREBP-1c and FAS (75, 76). Additionally, our results showed that liraglutide reduced the key regulators of de-novo lipogenesis, such as PPARG, SREBP-1C and LPL at the transcription level. This indicates that the decreased lipid accumulation is due to the inactivation of ACC which phosphorylated by p-AMPK and the decreased gene expression related to de-novo lipogenesis.

In conclusion, the changed gut microbiota diversity is associated with the altered SCFAs in chicken. SCFAs induce GLP-1 secretion in IECs via MAPK pathways. GLP-1 reduced hepatic fat synthesis by activating AMPK/ACC pathway. The result suggests that SCFAs-induced GLP-1 secretion links the regulation of gut microbiome on hepatic lipogenesis in chickens.

## Acknowledgement

This research was supported by earmarked fund for the National Key Research Program of China (2016YFD0500510), the China Agriculture Research System (CARS-40), and the Taishan Scholars Program (No.201511023).

## Declaration

The authors declare that they have no conflict of interests.

## Reference

1. Bäckhed F, Ding H, Wang T, Hooper LV, Koh GY, Nagy A, Semenkovich CF, Gordon JI. 2004. The gut microbiota as an environmental factor that regulates fat storage. Proc Natl Acad Sci U S A 101(44):15718–23. doi:10.1073/pnas.0407076101.PMID: 15505215.

2. Turnbaugh PJ, Ley RE, Mahowald MA, Magrini V, Mardis ER, Gordon JI. 2006. An obesity-associated gut microbiome with increased capacity for energy harvest. Nature 444(7122):1027–31. doi: 10.1038/nature05414. PMID: 17183312.

3. Membrez M, Blancher F, Jaquet M, Bibiloni R, Cani PD, Burcelin RG, Corthesy I, Macé K, Chou CJ. 2008. Gut microbiota modulation with norfloxacin and ampicillin enhances glucose tolerance in mice. FASEB J 22(7):2416–26. doi: 10.1096/fj.07-102723. PMID: 18326786.

4. Andersson U, Bränning C, Ahrné S, Molin G, Alenfall J, Onning G, Nyman M, Holm C. 2010. Probiotics lower plasma glucose in the high-fat fed C57BL/6J mouse. Benef Microbes 1(2):189–96. doi: 10.3920/BM2009.0036. PMID: 21840806

5. Tagliabue A, Elli M. 2013. The role of gut microbiota in human obesity: recent findings and future perspectives. Nutr Metab Cardiovasc Dis 23(3):160–8. doi: 10.1016/j.numecd.2012.09.002. PMID: 23149072.

6. Dahiya DK, Renuka, Puniya M, Shandilya UK, Dhewa T, Kumar N, Kumar S, Puniya AK, Shukla P. 2017. Gut Microbiota Modulation and Its Relationship with Obesity Using Prebiotic Fibers and Proboscis. Frontiers Microbiol 8:563. 10.3389/fmicb.2017.00563. PMID:28421057.

7. Rosenbaum M, Knight R, Leibel RL. 2015. The gut microbiota in human energy homeostasis and obesity. Trends Endocrinol Metab 26(9):493–501. doi: 10.1016/j.tem.2015.07.002. PMID: 26257300

8. Fuller R, Gibson GR. 1997. Modification of the intestinal microflora using probiotics and prebiotics. Scand J Gastroenterol Suppl 222:28-31. doi: 10.1080/00365521.1997.11720714. PMID: 9145443

9. Collins MD, Gibson GR. 1999. Probiotics, prebiotics, and synbiotics: approaches for modulating the microbial ecology of the gut. Am J Clin Nutr 69(5):1052S–1057S. doi: 10.1093/ajcn/69.5.1052s. PMID: 10232648.

10. Everard A, Cani PD. 2013. Diabetes, obesity and gut microbiota. Best Pract Res Clin Gastroenterol 27(1):73–83. doi: 10.1016/j.bpg.2013.03.007. PMID: 23768554.

11. Cremon C, Barbaro MR, Ventura M, Barbara G. 2018. Pre- and probiotic overview. Curr Opin Pharmacol 43:87–92. doi: 10.1016/j.coph.2018.08.010. PMID: 30219638.

12. Morrison DJ, Preston T. 2016. Formation of short chain fatty acids by the gut microbiota and their impact on human metabolism. Gut Microbes 7(3):189–200. doi: 10.1080/19490976.2015.1134082. PMID: 26963409.

13. Turnbaugh PJ, Ley RE, Mahowald MA, Magrini V, Mardis ER, Gordon JI. 2006. An obesity-associated gut microbiome with increased capacity for energy harvest. Nature 444(7122):1027–31. doi: 10.1038/nature05414. PMID: 17183312.

14. Schwiertz A, Taras D, Schäfer K, Beijer S, Bos NA, Donus C, Hardt PD. 2010. Microbiota and SCFA in lean and overweight healthy subjects. Obesity (Silver Spring) 18(1):190–5. doi: 10.1038/oby.2009.167. PMID:19498350.

15. Murphy EF, Cotter PD, Healy S, Marques TM, O’Sullivan O, Fouhy F, Clarke SF, O’Toole PW, Quigley EM, Stanton C, Ross PR, O’Doherty RM, Shanahan F. 2010. Composition and energy harvesting capacity of the gut microbiota: relationship to diet, obesity and time in mouse models.Gut 59(12):1635–42. doi: 10.1136/gut.2010.215665. PMID: 20926643.

16. Fernandes J, Su W, Rahat-Rozenbloom S, Wolever TM, Comelli EM. 2014. Adiposity, gut microbiota and faecal short chain fatty acids are linked in adult humans. Nutr Diabetes 4:e121. doi: 10.1038/nutd.2014.23. PMID: 24979150.

17. Hartstra AV, Bouter KE, Bäckhed F, Nieuwdorp M. 2015. Insights into the role of the microbiome in obesity and type 2 diabetes. Diabetes Care 38(1):159–65. doi: 10.2337/dc14-0769. PMID: 25538312.

18. Kasubuchi M, Hasegawa S, Hiramatsu T, Ichimura A, Kimura I. 2015. Dietary gut microbial metabolites, short-chain fatty acids, and host metabolic regulation. Nutrients 7(4):2839–49. doi: 10.3390/nu7042839. PMID: 25875123.

19. Gao Z, Yin J, Zhang J, Ward RE, Martin RJ, Lefevre M, Cefalu WT, Ye J. 2009. Butyrate improves insulin sensitivity and increases energy expenditure in mice. Diabetes 58(7):1509–17. doi: 10.2337/db08-1637. PMID: 19366864.

20. Lin HV, Frassetto A, Kowalik EJ Jr, Nawrocki AR, Lu MM, Kosinski JR, Hubert JA, Szeto D, Yao X, Forrest G, Marsh DJ. 2012. Butyrate and Propionate Protect against Diet-Induced Obesity and Regulate Gut Hormones via Free Fatty Acid Receptor 3-Independent Mechanisms. PLoS One 7(4):e35240. doi: 10.1371/journal.pone.0035240. PMID: 22506074.

21. Frost G, Sleeth ML, Sahuri-Arisoylu M, Lizarbe B, Cerdan S, Brody L, Anastasovska J, Ghourab S, Hankir M, Zhang S, Carling D, Swann JR, Gibson G, Viardot A, Morrison D, Louise Thomas E, Bell JD. 2014. The short-chain fatty acid acetate reduces appetite via a central homeostatic mechanism. Nat Commun 5:3611. doi: 10.1038/ncomms4611.PMID: 24781306.

22. den Besten G, Gerding A, van Dijk TH, Ciapaite J, Bleeker A, van Eunen K, Havinga R, Groen AK, Reijngoud DJ, Bakker BM. 2015. Protection against the metabolic syndrome by guar gum-derived short-chain fatty acids depends on peroxisome proliferator-activated receptor γ and glucagon-like peptide-1. PLoS One 10(8):e0136364. doi: 10.1371/journal.pone.0136364. PMID:26292284.

23. Lu Y, Fan C, Li P, Lu Y, Chang X, Qi K. 2016. Short chain fatty acids prevent high-fat-diet-induced obesity in mice by regulating G protein-coupled receptors and gut microbiota. Sci Rep 6:37589. doi: 10.1038/srep37589.PMID:27892486.

24. Covington DK, Briscoe CA, Brown AJ, Jayawickreme CK. 2006. The G protein coupled receptor 40 family (GPR40-43) and its role in nutrient sensing. Biochem Soc Trans 34(Pt 5):770–3. doi:10.1042/BST0340770. PMID:17052194.

25. Ge H, Li X, Weiszmann J, Wang P, Baribault H, Chen JL, Tian H, Li Y. 2008. Activation of G protein-coupled receptor 43 in adipocytes leads to inhibition of lipolysis and suppression of plasma free fatty acids. Endocrinology 149(9):4519–26. doi: 10.1210/en.2008-0059. PMID: 18499755.

26. Samuel BS, Shaito A, Motoike T, Rey FE, Backhed F, Manchester JK, Hammer RE, Williams SC, Crowley J, Yanagisawa M, Gordon JI. 2008. Effects of the gut microbiota on host adiposity are modulated by the short-chain fatty-acid binding G protein-coupled receptor, Gpr41. Proc Natl Acad Sci USA 105(43):16767–72. doi: 10.1073/pnas.0808567105. PMID: 18931303.

27. Zaibi MS, Stocker CJ, O’Dowd J, Davies A, Bellahcene M, Cawthorne MA, Brown AJ, Smith DM, Arch JR. 2010. Roles of GPR41 and GPR43 in leptin secretory responses of murine adipocytes to short chain fatty acids. FEBS Lett 584(11):2381–6. doi: 10.1016/j.febslet.2010.04.027. PMID: 20399779.

28. Kaji I, Karaki S, Kuwahara A. 2014. Short-chain fatty acid receptor and its contribution to glucagon-like peptide-1 release. Digestion 89(1):31–6. doi: 10.1159/000356211. PMID:24458110.

29. Tolhurst G, Heffron H, Lam YS, Parker HE, Habib AM, Diakogiannaki E, Cameron J, Grosse J, Reimann F, Gribble FM. 2012. Short-chain fatty acids stimulate glucagon-like peptide-1 secretion via the G-protein-coupled receptor FFAR2. Diabetes 61(2):364–71. doi: 10.2337/db11-1019. PMID: 22190648.

30. Psichas A, Sleeth ML, Murphy KG, Brooks L, Bewick GA, Hanyaloglu AC, Ghatei MA, Bloom SR, Frost G. 2015. The short chain fatty acid propionate stimulates GLP-1 and PYY secretion via free fatty acid receptor 2 in rodents. Int J Obes (Lond) 39(3):424–9. doi: 10.1038/ijo.2014.153. Epub 2014 Aug 11. PMID: 25109781.

31. Kimura I, Ozawa K, Inoue D, Imamura T, Kimura K, Maeda T, Terasawa K, Kashihara D, Hirano K, Tani T, Takahashi T, Miyauchi S, Shioi G, Inoue H, Tsujimoto G. 2013. The gut microbiota suppresses insulin-mediated fat accumulation via the short-chain fatty acidreceptor GPR43.Nat Commun 4:1829. doi: 10.1038/ncomms2852. PMID: 23652017

32. O’Hea EK, Leveille GA. 1969. Influence of fasting and refeeding on lipogenesis and enzymatic activity of pig adipose tissue. J Nutr 99(3):345-52. PMID: 5350990

33. Hermier D. 1997. Lipoprotein metabolism and fattening in poultry. Nutr 127(5 suppl): 805S–808S. doi: 10.1093/jn/127.5.805S. PMID: 9164241.

34. Mooney RA, Lane MD.1981. Formation and turn-over of triglyceride-rich vesicles in chick liver. Effects of cAMP and carnitine on triglyceride mobilization and conversion to ketones. J Biol Chem 256(22):11724-33. PMID: 6271759.

35. Van Deun K, Pasmans F, Ducatelle R, Flahou B, Vissenberg K, Martel A, Van den Broeck W, Van Immerseel F, Haesebrouck F. 2008. Colonization strategy of Campylobacter jejuni results in persistent infection of the chicken gut. Vet Microbiol 130(3-4):285–97. doi: 10.1016/j.vetmic.2007.11.027. PMID: 18187272.

36. Yuan C, He Q, Li JM, Azzam MM, Lu JJ, Zou XT. 2015. Evaluation of embryonic age and the effects of different proteases on the isolation and primary culture of chicken intestinal epithelial cells in vitro. Anim Sci J 86(6):588–94. doi: 10.1111/asj.12337. PMID: 2548860.

37. Kaiser A, Willer T, Sid H, Petersen H, Baumgärtner W, Steinberg P, Rautenschlein S.2016. Susceptibility of primary chicken intestinal epithelial cells for low pathogenic avian influenza virus and velogenic viscerotropic Newcastle disease virus. Virus Res 225:50–63. doi: 10.1016/j.virusres.2016.09.001. PMID: 2759673.

38. Kreymann B, Williams G, Ghatei MA, Bloom SR. 1987. Glucagon-like peptide-1 7-36: a physiological incretin in man. Lancet 2(8571):1300–4. PMID: 2890903.

39. Nishimura K, Hiramatsu K, Watanabe T, Kita K. 2017. Glucagon-like peptide-1 is co-localized with neurotensin in the chicken ileum. Cell Tissue Res 368(2):277–286. doi: 10.1007/s00441-016-2561-0. PMID: 28108848.

40. Hahn M, Gildemeister OS, Krauss GL, Pepe JA, Lambrecht RW, Donohue S, Bonkovsky HL. 1997. Effects of new anticonvulsant medications on porphyrin synthesis in cultured liver cells: potential implications for patients with acute porphyria. Neurology 49(1):97-106. PMID:9222176.

41. Su G, Crump D, Letcher RJ, Kennedy SW. 2014. Rapid in vitro metabolism of the flame retardant triphenyl phosphate and effects on cytotoxicity and mRNA expression in chicken embryonic hepatocytes. Environ Sci Technol 48(22):13511–9. doi: 10.1021/es5039547. PMID: 25350880.

42. Honda K. 2016. Glucagon-related peptides and the regulation of food intake in chickens. Anim Sci J. 87(9):1090–8. doi: 10.1111/asj.12619. PMID: 27150835.

43. Shen J, Obin MS, Zhao L. 2013. The gut microbiota, obesity and insulin resistance. Mol Aspects Med 34(1):39–58. doi: 10.1016/j.mam.2012.11.001. PMID: 23159341.

44. Caricilli AM, Saad MJ. 2014. Gut microbiota composition and its effects on obesity and insulin resistance. Curr Opin Clin Nutr Metab Care 17(4):312–8. doi: 10.1097/MCO.0000000000000067.PMID: 24848531.

45. Villanueva-Millán MJ, Pérez-Matute P, Oteo JA. 2015. Gut microbiota: a key player in health and disease. A review focused on obesity. J Physiol Biochem 71(3):509–25. doi: 10.1007/s13105-015-0390-3. Review.PMID: 2574993.

46. Geurts L, Lazarevic V, Derrien M, Everard A, Van Roye M, Knauf C, Valet P, Girard M, Muccioli GG, François P, de Vos WM, Schrenzel J, Delzenne NM, Cani PD. 2011. Altered gut microbiota and endocannabinoid system tone in obese and diabetic leptin-resistant mice: impact on apelin regulation in adipose tissue. Front Microbiol 2:149. doi: 10.3389/fmicb.2011.00149. PMID: 21808634.

47. Velasquez MT. 2018. Altered gut microbiota: a link between diet and the metabolic syndrome. Metab Syndr Relat Disord 16(7):321–328. doi: 10.1089/met.2017.0163. PMID: 29957105.

48. John GK, Mullin GE. 2016. The Gut Microbiome and Obesity. Curr Oncol Rep 18(7):45. doi: 10.1007/s11912-016-0528-7. PMID:27255389

49. Ley RE, Bäckhed F, Turnbaugh P, Lozupone CA, Knight RD, Gordon JI. 2005. Obesity alters gut microbial ecology. Proc Natl Acad Sci USA 102(31):11070–5. doi: 10.1073/pnas.0504978102. PMID: 16033867.

50. Macfarlane S, Macfarlane GT. 2003. Regulation of short-chain fatty acid production. Proc Nutr Soc 62(1):67-72. 10.1079/PNS2002207. PMID:12740060.

51. Pessione E. 2012. Lactic acid bacteria contribution to gut microbiota complexity: lights and shadows. Front Cell Infect Microbiol 2:86. doi: 10.3389/fcimb.2012.00086. PMID: 22919677.

52. Meimandipour A, Hair-Bejo M, Shuhaimi M, Azhar K, Soleimani AF, Rasti B, Yazid AM. 2010. Gastrointestinal tract morphological alteration by unpleasant physical treatment and modulating role of Lactobacillus in broilers. Br Poult Sci. Br Poult Sci 51(1):52–9. doi: 10.1080/00071660903394455. PMID: 20390569.

53. Liang Y, Lin C, Zhang Y, Deng Y, Liu C, Yang Q. 2018. Probiotic mixture of Lactobacillus and Bifidobacterium alleviates systemic adiposity and inflammation in non-alcoholic fatty liver disease rats through Gpr109a and the commensal metabolite butyrate. Inflammopharmacology 26(4):1051–1055. doi: 10.1007/s10787-018-0479-8. PMID:29633106.

54. Wang H, Ni X, Qing X, Zeng D, Luo M, Liu L, Li G, Pan K, Jing B. 2017. Live Probiotic Lactobacillus johnsonii BS15 Promotes Growth Performance and Lowers Fat Deposition by Improving Lipid Metabolism, Intestinal Development, and Gut Microflora in Broilers.Front Microbiol 8:1073. doi: 10.3389/fmicb.2017.01073.PMID:28659893

55. Tamura M, Hirayama K, Itoh K. 1999. Effects of guar gum and cellulose on cecal enzyme activity and cecal short-chain fatty acids in young and aged mice. Ann NutrMetab 43(1):60–5.doi:10.1159/000012768. PMID:10364632.

56. Butt MS, Shahzadi N, Sharif MK, Nasir M. 2007. Guar gum: a miracle therapy for hypercholesterolemia, hyperglycemia and obesity. Crit Rev Food Sci Nutr 47(4):389–96. doi: 10.1080/10408390600846267. PMID: 17457723.

57. Krotkiewski M. 1984. Effect of guar gum on body-weight, hunger ratings and metabolism in obese subjects. Br J Nutr. Jul 52(1):97-105. doi. 10.1079/BJN19840075PMID: 6331498.

58. Chambers ES, Viardot A, Psichas A, Morrison DJ, Murphy KG, Zac-Varghese SE, MacDougall K, Preston T, Tedford C, Finlayson GS, Blundell JE, Bell JD, Thomas EL, Mt-Isa S, Ashby D, Gibson GR, Kolida S, Dhillo WS, Bloom SR, Morley W, Clegg S, Frost G. 2015. Effects of targeted delivery of propionate to the human colon on appetite regulation, body weight maintenance and adiposity in overweight adults. Gut 64(11):1744–54. doi: 10.1136/gutjnl-2014-307913. PMID: 25500202.

59. Heimann E, Nyman M, Degerman E. 2014. Propionic acid and butyric acid inhibit lipolysis and de novo lipogenesis and increase insulin-stimulated glucose uptake in primary rat adipocytes. Adipocyte 4(2):81–8. doi: 10.4161/21623945.2014.960694. PMID: 26167409.

60. Hudson BD, Tikhonova IG, Pandey SK, Ulven T, Milligan G. 2012. Extracellular ionic locks determine variation in constitutive activity and ligand potency between species orthologs of the free fatty acid receptors FFA2 and FFA3. J Biol Chem 287(49):41195–209. doi: 10.1074/jbc.M112.396259. PMID: 23066016

61. Christiansen CB, Gabe MBN, Svendsen B, Dragsted LO, Rosenkilde MM, Holst JJ. 2018. The impact of short-chain fatty acids on GLP-1 and PYY secretion from the isolated perfused rat colon. Am J Physiol Gastrointest Liver Physiol 315(1):G53–G65. doi: 10.1152/ajpgi.00346.. PMID:29494208.

62. Brown AJ, Goldsworthy SM, Barnes AA, Eilert MM, Tcheang L, Daniels D, Muir AI, Wigglesworth MJ, Kinghorn I, Fraser NJ, Pike NB, Strum JC, Steplewski KM, Murdock PR, Holder JC, Marshall FH, Szekeres PG, Wilson S, Ignar DM, Foord SM, Wise A, Dowell SJ. 2003. The Orphan G protein-coupled receptors GPR41 and GPR43 are activated by propionate and other short chain carboxylic acids. J Biol Chem 278(13):11312-9.PMID:12496283.

63. Le Poul E, Loison C, Struyf S, Springael JY, Lannoy V, Decobecq ME, Brezillon S, Dupriez V, Vassart G, Van Damme J, Parmentier M, Detheux M. 2003. Functional characterization of human receptors for short chain fatty acids and their role in polymorphonuclear cell activation. J Biol Chem 278:25481–25489. doi: 10.1074/jbc.M301403200. PMID: 12711604.

64. Yonezawa T, Haga S, Kobayashi Y, Katoh K, Obara Y. 2008. Unsaturated fatty acids promote proliferation via ERK1/2 and Akt pathway in bovine mammary epithelial cells. Biochem Biophys Res Commun 367(4):729–35. doi: 10.1016/j.bbrc.2007.12.190. PMID:18191634

65. Seljeset S, Siehler S. 2012. Receptor-specific regulation of ERK1/2 activation by members of the “free fatty acid receptor” family. J Recept Signal Transduct Res 32(4):196–201. doi: 10.3109/10799893.2012.692118. PMID: 22712802.

66. Yonezawa T, Haga S, Kobayashi Y, Katoh K, Obara Y. 2009. Short-chain fatty acid signaling pathways in bovine mammary epithelial cells. Regul Pept 153(1-3):30–6. doi: 10.1016/j.regpep.2008.11.012. PMID: 19101595.

67. Huang G, Shi LZ, Chi H. 2009. Regulation of JNK and p38 MAPK in the immune system: signal integration, propagation and termination. Cytokine 48(3):161–9. doi: 10.1016/j.cyto.2009.08.002. PMID:19740675

68. Blaut M. 2015. Gut microbiota and energy balance: role in obesity. Proc Nutr Soc 74(3):227–34. doi: 10.1017/S0029665114001700. PMID:25518735.

69. Vilsbøll T, Christensen M, Junker AE, Knop FK, Gluud LL. 2012. Effects of glucagon-like peptide-1 receptor agonists on weight loss: systematic review and meta-analyses of randomised controlled trials. BMJ 344:d7771. doi: 10.1136/bmj.d7771. PMID: 22236411.

70. Petit JM, Vergès B. 2017. GLP-1 receptor agonists in NAFLD. Diabetes Metab 43 Suppl 1:2S28–2S33. doi: 10.1016/S1262-3636(17)30070-8. PMID: 28431668.

71. Lu Y, Fan C, Li P, Lu Y, Chang X, Qi K. 2016. Short Chain Fatty Acids Prevent High-fat-diet-induced Obesity in Mice by Regulating G Protein-coupled Receptors and Gut Microbiota.Sci Rep 6:37589. doi: 10.1038/srep37589.PMID:27892486.

72. Ding X, Saxena NK, Lin S, Gupta NA, Anania FA. 2006. Exendin-4, a glucagon-Like protein-1 (GLP-1) receptor agonist, reverses hepatic steatosis in ob/ob mice. Hepatology 43(1):173–81. PMID: 16374859.

73. Hardie DG, Pan DA. 2002. Regulation of fatty acid synthesis and oxidation by the AMP-activated protein kinase. Biochem Soc Trans 30(Pt 6):1064–70. doi:10.1042/. PMID: 12440973

74. Ben-Shlomo S, Zvibel I, Shnell M, Shlomai A, Chepurko E, Halpern Z, Barzilai N, Oren R, Fishman S. 2011. Glucagon-like peptide-1 reduces hepatic lipogenesis via activation of AMP-activated protein kinase. J Hepatol 54(6):1214–23. doi: 10.1016/j.jhep.2010.09.032. PMID: 21145820.

75. Viollet B, Foretz M, Guigas B, Horman S, Dentin R, Bertrand L, Hue L, Andreelli F. 2006. Activation of AMP-activated protein kinase in the liver: a new strategy for the management of metabolic hepatic disorders. J Physiol 574(Pt 1):41-53. doi: 10.1113/jphysiol.2006.108506. PMID: 16644802.

76. Postic C, Girard J. 2008. Contribution of de novo fatty acid synthesis to hepatic steatosis and insulin resistance: lessons from genetically engineered mice. J Clin Invest. 118(3):829–38. doi: 10.1172/JCI34275. PMID: 18317565.

